# Heart rate variability impairment during sleep in Veterans with REM sleep behavior disorder, traumatic brain injury, and posttraumatic stress disorder: An early potential window into autonomic dysfunction?

**DOI:** 10.1101/2024.09.20.614142

**Authors:** Hannah A. Cunningham, Laura Dovek, Natasha Recoder, Mohini D. Bryant-Ekstrand, Brittany R. Ligman, Juan Piantino, Miranda M. Lim, Jonathan E. Elliott

**Affiliations:** Oregon Health & Science University, Department of Behavioral Neuroscience, Portland, OR, USA; VA Portland Health Care System, Research Service, Portland, OR, USA; Oregon Health & Science University, Department of Neurology, Portland, OR, USA; Oregon Health & Science University, Department of Pediatrics, Portland, OR, USA; VA Portland Health Care System, VISN 20 Northwest Mental Illness Research, Education and Clinical Center (MIRECC), Portland, OR, USA; VA Portland Health Care System, National Center for Rehabilitative Auditory Research (NCRAR), and Neurology, Portland, OR, USA; Oregon Health & Science University, Department of Medicine; Division of Pulmonary and Critical Care Medicine and Oregon Institute of Occupational Health Sciences, Portland, OR, USA

**Author notes:** **Corresponding author and address for reprints.** Jonathan E. Elliott, PhD, VA Portland Health Care System, 3710 SW US Veterans Hospital Road, Mailcode P3-RD42, Portland, OR 97239. Dr. Elliott takes full responsibility for the data, the analyses and interpretation, and the conduct of the research; has full access to all of the data; and has the right to publish any and all data separate and apart from any sponsor. **Address Where This Work Was Conducted.** VA Portland Health Care System, 3710 SW US Veterans Hospital Road, Mail code P3-RD42, Portland, OR 97239.

**Keywords:** REM sleep behavior disorder, heart rate variability, neurotrauma, PTSD, traumatic brain injury

## Abstract

Individuals with comorbid REM sleep behavior disorder (RBD) and neurotrauma (defined by traumatic brain injury and post-traumatic stress disorder) have an earlier age of RBD symptom onset, increased RBD-related symptom severity and more neurological features indicative of prodromal synucleinopathy compared to RBD only. An early sign of neurodegenerative condition is autonomic dysfunction, which we sought to evaluate by examining heart rate variability during sleep. Participants with overnight polysomnography were recruited from the VA Portland Health Care System. Veterans without neurotrauma or RBD (controls; n=19), with RBD only (RBD, n=14), and with RBD and neurotrauma (RBD+NT, n=19) were evaluated. Eligible 5-minute NREM and REM epochs without apneas/hypopneas, microarousals, and ectopic beats were analyzed for frequency and time domain (e.g. low frequency power, LF; high frequency power, HF; root mean square of successive RR intervals, RMSSD; % of RR intervals that vary ≥50 ms, pNN50) heart rate variability outcomes. Heart rate did not significantly differ between groups in any sleep stage. Time domain and frequency domain variables (e.g., LF power, HF power, RMSSD, and pNN50) were significantly reduced in the RBD and RBD+NT groups compared to controls and RBD only during NREM sleep. There were no group differences detected during REM sleep. These data suggest significant reductions in heart rate variability during NREM sleep in RBD+NT participants, suggesting greater autonomic dysfunction compared to controls or RBD alone. Heart rate variability during sleep may be an early, promising biomarker, yielding mechanistic insight for diagnosis and prognosis of early neurodegeneration in this vulnerable population.

**STATEMENT OF SIGNIFICANCE:** Comorbid REM sleep behavior disorder (RBD) and neurotrauma (NT, traumatic brain injury + post-traumatic stress disorder; RBD+NT) is associated with increased neurodegenerative symptom burden and worsened health. Sleep and autonomic function are integrally and bidirectionally related to neurodegenerative processes. In the current study, we sought to determine if early signs of autonomic dysfunction, measured via heart rate variability (HRV), were present during sleep in comorbid RBD+NT compared to RBD only and controls. Our data show reduced time and frequency domain HRV during NREM sleep in RBD+NT Veterans compared to RBD only and controls. These data contribute evidence that participants with RBD and comorbid NT demonstrate significantly worse autonomic dysfunction compared to age/sex matched participants with RBD alone.

## INTRODUCTION

Idiopathic rapid eye movement (REM) sleep behavior disorder (RBD) is a sleep disorder characterized by the loss of skeletal muscle atonia during REM sleep. RBD is regarded as a prodromal synucleinopathy; 40%-70% of patients will phenoconvert to overt synucleinopathy (e.g. Parkinson’s disease (PD), dementia with Lewy bodies (DLB), or multiple system atrophy (MSA)) within 5-10 years [1-3]. A recent systematic review and meta-analysis suggest that individuals with RBD have a 96% chance of developing overt synucleinopathy within 14 years [4]. Many neurological conditions, including Alzheimer’s [5], DLB [6], and PD [7, 8], have impaired autonomic function determined via heart rate variability (HRV). HRV is a measure of cardiac autonomic modulation via parasympathetic and sympathetic pathways and is an established marker of cardiac health and cardiovascular risk, strongly correlating with overall morbidity and mortality [9-12]. During wake, RBD patients have general autonomic dysfunction [13], though not all studies observed a change in wake HRV [14]. However, HRV during sleep may reveal critical differences; in contrast to individuals without RBD, patients with RBD surprisingly did not differ in spectral HRV in NREM versus REM sleep [15]. Additionally, patients with RBD have lower frequency domain and time domain variables of HRV [16, 17].

Comorbid neurotrauma (NT) is defined by a combination of traumatic brain injury (TBI) and post-traumatic stress disorder (PTSD). Similar to RBD, comorbid neurotrauma is a risk factor for neurodegenerative disorders like PD and dementia [18]. While many studies have examined HRV changes with TBI and PTSD alone, none have examined the comorbid condition. The comorbid condition of TBI+PTSD has been proposed as a worthwhile and distinct entity for research and clinical management [19-21]. HRV in patients with PTSD is lower during wake [22-24], negatively correlated with worse PTSD symptoms [25], and predicts treatment outcomes [26]. TBI is also associated with lower HRV during wake[27].

Cross sectional data from our laboratory has shown that participants with both RBD and NT (i.e., RBD+NT) have earlier age of RBD symptom onset, increased symptom severity and generally worsened health and neurological function compared to individuals with only RBD [21]. This indicates a need to distinguish between those with RBD+NT from RBD alone. It is unknown how NT and RBD influences overnight HRV. Therefore, in the current study we examine HRV during overnight PSG for patients with RBD and patients with RBD+NT compared to patients with neither condition. We hypothesize that HRV will be reduced during sleep in Veterans with RBD+NT compared to age- and sex-matched Veterans with RBD only and control participants with neither RBD nor NT.

## METHODS

### Study type

This is a retrospective, case-control analysis of data collected from a single site across two studies. All participants underwent overnight polysomnography in an AASM-accredited sleep laboratory (see details below). Participants were selected consecutively, matched by age (+/-4 years) and sex, for analysis according to prespecified criteria. Cases included 3 groups: 1) RBD+NT (i.e., co-morbid RBD, TBI and PTSD), 2) RBD only (i.e., RBD without NT), and 3) controls without RBD or NT. PSG studies had to be diagnostic (i.e. no split night), AHI < 25, and time in bed greater than four hours.

### Participant Enrollment

All subjects were recruited from the VA Portland Health Care System between May 2015 and April 2024 as part of the North American Prodromal Synucleinopathy (NAPS) consortium [28] or as part of a single site, prospective cross-sectional study (previously described in [29-31]). Participants were included if they met diagnostic criteria, including a diagnostic study and PSG recording time longer than four hours. N=419 studies were considered for analysis; 362 were excluded either for not meeting PSG inclusion criteria or group inclusion criteria. Of the remaining, two were excluded due to no eligible 5-minute interval and three for an irregular heart rhythm. The remaining 52 participants were included in the final analysis (n=19 as controls, n=14 with RBD only, and n=19 with RBD+NT). The VA Portland Health Care System Institutional Review Board approved the studies (IRB #4542 and IRB #3641), and the studies were performed according to principles of The Belmont Report and the *Declaration of Helsinki*.

Participants reported to the sleep clinic prior to their usual bedtime. A sleep technician fitted the participants with 12-lead polysomnography, 3-lead electrocardiogram, electromyogram to assess muscle tone, and a nasal canula used to assess apneas and hypopneas. All sleep data were acquired and staged according to the American Academy of Sleep Medicine (AASM) standards using NihonKohden Polysmith software.

### Clinical Diagnoses

RBD was diagnosed according to International Classification of Sleep Disorders (ICSD)-3 clinical criteria, including presence of REM sleep without atonia on video-polysomnography coupled with a history of dream enactment behavior. NT was assessed using the Head Trauma Events Characteristics (HTEC) for TBI in conjunction with standard cutoff criteria using the PTSD Checklist for DSM-5 (PCL-5) for PTSD as previously described [21, 30]. All participants in the RBD+NT group met diagnostic criteria for both PTSD and TBI. Participants in the control and RBD only group did not meet criteria for TBI or PTSD.

### Sleep Staging and Heart Rate Variability Analysis

American Academy of Sleep Medicine-accredited polysomnographic technicians scored the sleep staging for 30 second epochs for all overnight sleep studies. Each epoch was scored as wake, stage 1 (N1), stage 2 (N2), stage 3 (N3), and REM sleep. For heart rate variability analysis, a single trained staff member identified 5-minute epochs that were free of apneas, hypopneas, microarousals, ectopic beats, and changes in sleep stage in the epoch immediately prior and after. Eligible epochs were collapsed into NREM and REM. Stage N1 sleep was not used for analysis.

WinCPRS (Absolute Aliens, Turku Finland) was used to conduct analyses of HRV to calculate time domain and frequency domain variables. The following HRV variables were anlayzed: low frequency power (LF), high frequency power (HF), total power, the ratio of LF and HF (LF/HF), root mean square of successive normal RR intervals (RMSSD), the percentage of successive normal RR intervals that vary by 50ms (pNN50) and the standard deviation of successive normal RR intervals (SDNN). R waves were automatically marked, and a trained research assistant manually confirmed all R waves were properly marked. All HRV variables were calculated for all 5-minutes epochs identified for every participant. If a participant had multiple 5-minute epochs for the same sleep stage, the data were averaged to produce a single value for the stage. NREM stages of N2 and N3 were averaged into a single stage, “NREM”. Of the 52 participants enrolled, all had an eligible NREM interval. One RBD participant and two RBD+NT participants did not have any eligible REM epochs. Wake was not analyzed due to A) limited resting wake time prior to “lights out” because the overnight PSG studies were conducted in a clinical setting; and B) high inter-individual variability in stable wake intervals (e.g., no movement or undisturbed) throughout the night.

### Statistical Analysis

Statistical analyses were performed using R Studio Version 4.3.2 and GraphPad Prism 10.2.3 and applied a significance level of *P <*0.05. Data are presented as either mean ± standard deviation or natural logarithmic mean ± standard deviation, unless otherwise specified. Using the Shapiro-Wilk test, all participant demographic data was tested for normal distribution. For variables in which all cohorts passed the normality test (NREM, percentage and minutes; N2 sleep, percentage and minutes; REM sleep, percentage and minutes) an Ordinary one-way analysis of variance (ANOVA) was run. Otherwise, Kruskal-Wallis test was used. We used Shapiro-Wilks test to assess normality and natural logarithmic transformed the non-normal variables (e.g., RMSSD, SDNN, LF, HF, LF/HF, total power). To examine group differences, we used an Ordinary one-way ANOVA with a pairwise t-test. pNN50 continued to be non-normally distributed after natural log transformation, so we proceeded with Kruskal-Wallis rank sum test analyses on the non-transformed values with a pairwise Wilcoxon signed-rank test.

## RESULTS

Participant characteristics are presented in **Table 1**. The groups were age- and sex-matched. There were no significant differences in sleep characteristics.

**Table 1.** Participant demographic and sleep characteristics.

|  | **Controls**<br>**n=19** | **RBD**<br>**n=14** | **RBD + NT**<br>**n=19** | **Test**<br>**Statistic** | **P-value** |
| --- | --- | --- | --- | --- | --- |
| Age, years | 51.95 ± 14.4 | 50.93 ± 19.9 | 51.05 ± 15.0 | 0.1 | 0.964 |
| Sex, male (%) | 17 (89.5%) | 13 (92.9%) | 18 (94.7%) | 0.4 | 0.831 |
| Race (%) |  |  |  |  |  |
| White | 18 (94.74%) | 12 (85.7%) | 13 (68.4%) | 4.6 | 0.099 |
| Black or African | 0 (0%) | 0 (0%) | 1 (5.2%) | 1.7 | 0.420 |
| American<br>Indian/Alaska Native | 1 (5.26%) | 0 (0%) | 0 (0%) | 1.7 | 0.420 |
| Unknown/ Not<br>Reported | 0 (0%) | 2 (14.3%) | 5 (25%) | 5.6 | 0.062 |
| TST, hrs | 7.81 ± 8.2 | 5.94 ± 0.9 | 5.38 ± 1.3 | 3.0 | 0.228 |
| WASO, min | 77.26 ± 60.0 | 42.57 ± 30.7 | 78.82 ± 55.9 | 6.0 | 0.050* |
| Sleep Efficiency, % | 79.66 ± 12.6 | 83.96 ± 11.9 | 74.66 ± 17.6 | 3.9 | 0.142 |
| Sleep Latency, min | 15.82 ± 17.1 | 27.11 ± 27.3 | 29.71 ± 29.5 | 2.5 | 0.283 |
| N1 Sleep, % | 11.66 ± 6.7 | 9.87 ± 6.4 | 15.14 ± 9.92 | 3.0 | 0.223 |
| N1 Sleep, min | 42.21 ± 26.6 | 32.36 ± 14.0 | 44.18 ± 25.6 | 0.9 | 0.627 |
| N2 Sleep <sup>a</sup>, % | 68.09 ± 11.6 | 65.90 ± 8.1 | 65.02 ± 12.7 | 0.4 | 0.691 |
| N2 Sleep <sup>a</sup>, min | 243.16 ± 57.1 | 235.29 ± 51.0 | 213.18 ± 75.9 | 1.1 | 0.331 |
| N3 Sleep, % | 4.16 ± 9.1 | 7.26 ± 7.7 | 5.06 ± 7.6 | 2.9 | 0.235 |
| N3 Sleep, min | 14.16 ± 34.0 | 25.96 ± 26.3 | 16.18 ± 23.4 | 3.6 | 0.166 |
| NREM Sleep <sup>a</sup>, % | 83.91 ± 6.4 | 83.03 ± 7.1 | 85.22 ± 9.7 | 0.3 | 0.727 |
| NREM Sleep <sup>a</sup>, min | 299.53 ± 49.9 | 293.61 ± 40.6 | 273.55 ± 70.8 | 1.1 | 0.346 |
| REM Sleep <sup>a</sup>, % | 16.07 ± 6.4 | 16.99 ± 7.1 | 14.76 ± 9.7 | 0.3 | 0.720 |
| REM Sleep <sup>a</sup>, min | 58.71 ± 27.5 | 62.50 ± 28.6 | 49.26 ± 33.2 | 0.9 | 0.420 |
| AHI | 7.82 ± 8.6 | 6.36 ± 4.2 | 8.27 ± 9.3 | 0.1 | 0.955 |

<sup>a</sup> Data set passed Shapiro Wilk test for Normality and an Ordinary One-Way ANOVA was performed. Otherwise, comparisons performed were Kruskal-Wallis test.
Values are mean ± SD or count (%); AHI, apnea hypopnea index; NREM, non-REM sleep; REM, rapid eye movement; TST, total sleep time; WASO, wake after sleep onset.

The stage specific results for REM and NREM are tabulated in **Table 2**. There were no significant differences in heart rate between the groups in any sleep stage (REM p=0.672; NREM p=0.356). There were significant group differences in frequency domain variables. High frequency (HF) power during NREM was significantly different, (p=0.017), whereby the RBD+NT group (4.5±1.4 a.u.) was significantly lower compared to RBD (5.9±1.5 a.u., p=0.009) and controls (5.7±1.7 a.u., p=0.022). Additionally, low frequency (LF) power was reduced during NREM (p=0.010) in the RBD+NT group (5.4±1.5 a.u.) compared to RBD (6.1±2.0 a.u.; p=0.006) and controls (6.5±1.5 a.u.; p=0.025). Group differences for LF and HF power are graphed in **Figure 1a** and **1b**. Total power (p=0.023) was reduced during NREM (RBD+NT, 6.3±1.1 a.u. vs. RBD 7.5±1.3 a.u., p=0.013; and RBD+NT vs controls 7.2±1.3, a.u., p=0.031). There were no significant group differences in REM for LF, HF or total power. Additionally, there was no significant group effect for LF/HF ratio (p=0.511) during NREM and REM.

**Table 2.** Nocturnal HRV Characteristics by Sleep Stage.

|  | Controls | RBD | RBD+NT | p-value<sup>a</sup> |
| --- | --- | --- | --- | --- |
| **NREM** |  |  |  |  |
| N | 19 | 14 | 19 |  |
| HR, beats/min | 65 ± 10 | 62 ± 12 | 67 ± 11 | 0.356 |
| IBI, ms | 952 ± 139 | 1,012 ± 191 | 913 ± 137 | 0.203 |
| SDNN, ms | 3.7 ± 0.7 | 3.8 ± 0.6 | 3.3 ± 0.5<sup>b</sup> | 0.023* |
| RMSSD, ms | 3.6 ± 0.9 | 3.7 ± 0.8 | 3.0 ± 0.7<sup>b</sup> | 0.024* |
| pNN50, %<sup>†</sup> | 5.0 (1.1, 15.6) | 9.2 (1.2, 33.6) | 0.3 (0.0, 9.0)<sup>b</sup> | 0.026* |
| Total power, ms<sup>2</sup> | 7.2 ± 1.3 | 7.5 ± 1.3 | 6.3 ± 1.1<sup>b</sup> | 0.023* |
| LF, ms<sup>2</sup> | 6.2 ± 1.4 | 6.6 ± 1.4 | 5.2 ± 1.2<sup>b</sup> | 0.010* |
| HF, ms<sup>2</sup> | 5.7 ± 1.7 | 5.9 ± 1.5 | 4.5 ± 1.4<sup>b</sup> | 0.017* |
| LF/HF, a.u. | 5.2 ± 0.9 | 5.4 ± 0.8 | 5.5 ± 1.0 | 0.511 |
| **REM** |  |  |  |  |
| N | 19 | 13 | 17 |  |
| HR, beats/min | 66 ± 9 | 63 ± 12 | 66 ± 10 | 0.672 |
| IBI, ms | 933 ± 113 | 984 ± 175 | 934 ± 132 | 0.539 |
| SDNN<sub>ln</sub>, a.u. | 4.0 ± 0.6 | 3.8 ± 0.7 | 3.7 ± 0.7 | 0.313 |
| RMSSD<sub>ln</sub>, a.u. | 3.5 ± 0.9 | 3.5 ± 0.8 | 2.9 ± 0.7 | 0.081 |
| pNN50, %<sup>†</sup> | 3.9 (1.8, 14.6) | 8.5 (0.4, 27.1) | 1.4 (0.0, 7.6) | 0.163 |
| Total power<sub>ln</sub>, a.u. | 7.8 ± 1.3 | 7.5 ± 1.5 | 7.0 ± 1.5 | 0.236 |
| LF<sub>ln</sub>, a.u. | 6.5 ± 1.5 | 6.1 ± 2.0 | 5.4 ± 1.5 | 0.164 |
| HF<sub>ln</sub>, a.u. | 5.3 ± 1.9 | 5.3 ± 1.6 | 4.1 ± 1.6 | 0.082 |
| LF/HF<sub>ln</sub>, a.u. | 5.9 ± 1.0 | 5.5 ± 1.2 | 6.0 ± 1.0 | 0.436 |

<sup>a</sup> Analysis of Variance test used unless otherwise indicated.
<sup>†</sup> Kruskal-Wallis rank sum test used.
<sup>b</sup> RBD+NT < Controls and RBD
Values are mean ± SD or median (Q2, Q3); HF<sub>ln</sub>, natural log transformed high frequency-HRV; HR, heart rate; HRV, heart rate variability; IBI, interbeat interval; LF<sub>ln</sub>, natural log transformed low frequency-HRV; LF/HF<sub>ln</sub>, natural log transformed low frequency/high frequency HRV ratio; NREM, non-rapid eye movement sleep; pNN50, percentage of R-R intervals that varied by 50 ms or more; REM, rapid eye movement sleep; RMSSD<sub>ln</sub>, natural log transformed root mean squared of successive differences of R-R intervals; SDNN<sub>ln</sub>, natural log transformed standard deviation of NN intervals.

**Figure 1.**
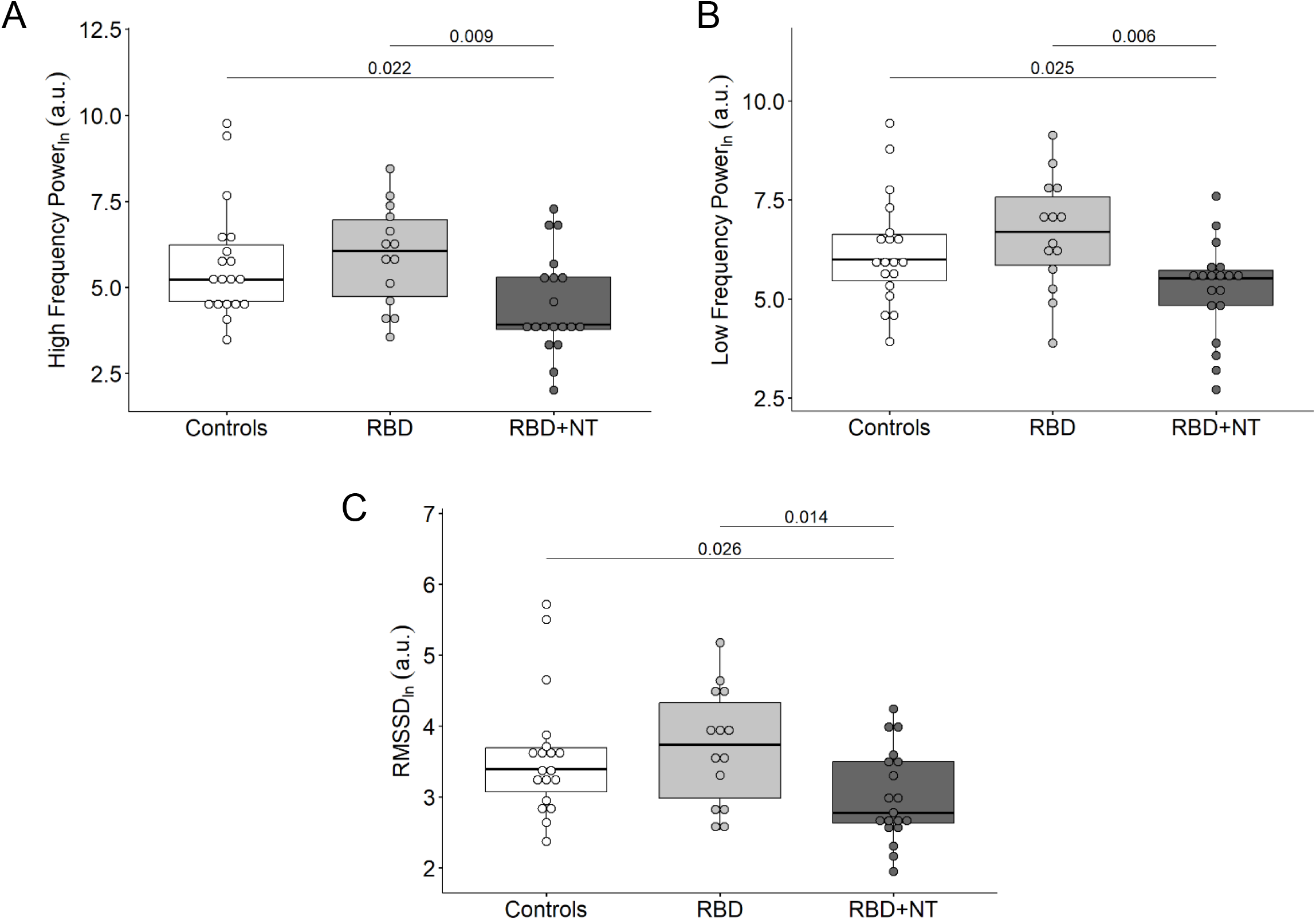
Group differences during NREM. The RBD+NT group has significantly reduced high frequency (A), low frequency power (B), and RMSSD (C) values compared to controls and RBD only. pairwise t-test values are noted on the graphs. Boxes are interquartile range (IQR). Whiskers represent max and min value no further than 1.5 * IQR. Variables are natural logarithmic transformed. RBD, REM sleep behavior disorder; RMSSD, root mean square of successive differences of RR intervals.

Group effects were significant for RMSSD (p=0.024) during NREM (RBD+NT, 3.0±0.7 a.u. vs RBD, 3.7±0.8 a.u., p=0.015; and RBD+NT vs controls, 3.6±0.9 a.u., p=0.04). These differences are graphed in **Figure 1c**. Similarly, pNN50 (p=0.026) was reduced during NREM (RBD+NT, 0.3 [0.0, 9.0]% vs. RBD 9.2 [1.2, 33.6]%, p=0.015; and RBD+NT vs controls 5.0 [1.1, 15.6]%, p=0.04; median [25% quartile, 75% quartile]). SDNN (p=0. 0297) was reduced during NREM (RBD+NT, 3.3±0.5 ms vs. RBD 3.8±0.6 ms, p=0.014; and RBD+NT vs controls 3.7±0.7, ms, p=0.044). There were no significant differences during REM for RMSSD, pNN50, and SDNN.

## DISCUSSION

Comorbid REM sleep behavior disorder (RBD) and neurotrauma (NT, traumatic brain injury, TBI + post-traumatic stress disorder, PTSD) is a distinct clinical group that has earlier onset of RBD symptoms and more neurological impairment concerning prodromal synucleinopathies [21, 29]. Our present study sought to examine heart rate variability (HRV) during sleep in participants with RBD+NT compared to participants with RBD only and to controls without RBD or NT. Our groups were age- and sex-matched without statistical differences in any sleep stage amount (minutes or percentage). The present analysis demonstrates significant reductions in HRV (e.g., LF power, HF power, RMSSD, and pNN50) during NREM sleep in Veterans with RBD+NT compared to age- and sex-matched participants with RBD only and controls without RBD or NT. These data may provide a potential indicator of autonomic dysfunction in this clinical population.

Work by others has shown patients with RBD have lower HRV in both frequency and time domain variables [16, 17], though not all studies observe a change in HRV during wake [18]. Lanfranchi et al., observed patients with RBD do not exhibit the hallmark differences in HRV between NREM and REM sleep [15]. In controls, they observed reduced HF components during REM sleep, indicative of parasympathetic withdrawal. However, the RBD participants had blunted parasympathetic withdrawal during REM sleep and no differences in NREM sleep measures. This is in contrast to our study which found no differences between the RBD and control groups, i.e., statistical differences were confined to the RBD+NT group. This discrepancy may be a result of several differences, including participant age and our explicit exclusion of NT in the RBD only group. On average participants with RBD (RBD only or RBD+NT) were 51 years of age compared to 60 years of age in Lanfranchi et al [15]. It is unknown whether autonomic dysfunction progressively declines with disease duration, and its relationship to risk of phenoconversion is contested [32-34].

The parasympathetic nervous system is dominate during NREM sleep and physiological signals (e.g., heart rate, blood pressure) are characteristically rhythmic and stable [35-37]. In contrast, sympathetic nervous system activity dominates during REM sleep compared to parasympathetic activity, resulting in a highly variable heart rate, respiratory rate, and blood pressure response. Consequently, we can speculate our finding could reflect dysfunction of autonomic control specific to NREM sleep. Studies administering atropine show the HF power and portions of the LF power are associated with parasympathetic nervous system activity [38]. We observed reduction in both bands during NREM sleep. Our results could reflect general parasympathetic dysfunction; however, this is speculative and would need to be experimentally tested with modalities that directly measure parasympathetic or sympathetic activity during sleep (e.g., microneurography of muscle sympathetic nerve activity). Alternatively, the inherent stability of these signals during NREM sleep—in comparison to the volatility of these signals during REM sleep—may provide a more accurate description of general autonomic function. Alterations to HRV could reflect an early symptom of autonomic dysfunction. Neurological conditions, including Parkinson’s disease, have altered autonomic activity early in the disease course [39-41]. Dysautonomia is one of the first non-motor symptoms of Parkinson’s disease and can precede diagnosis by at least 5 years but is estimated to be 11 to 20 years before phenoconversion [42]. With 40%-70% of RBD patients phenoconverting to overt synucleinopathy within 5-10 years [1-3], alteration to HRV as measured during NREM sleep could be a prodromal symptom of developing autonomic dysfunction. Indeed, patients with RBD display general autonomic dysfunction [21, 32, 43]. Given the higher prevalence of RBD in Veterans with NT and the earlier age of RBD symptom onset [29], further longitudinal research is needed to confirm the progression of autonomic dysfunction throughout the disease course.

Although also speculative, the changes we report in autonomic function during NREM sleep in this potentially vulnerable/at risk population may reflect more widespread changes in other NREM sleep dominant systems, including glymphatic function (for review: [44]). The recently described glymphatic system, is a brain-wide network of perivascular spaces that promotes interstitial solute clearance and is preferentially active during slow-wave sleep with potential autonomic regulation [45-48] and has been shown to be impaired following TBI [49-51]. Additional work is needed to confirm whether RBD alone contributes to glymphatic impairment, though previous studies suggest a possible reduction in glymphatic function in RBD [52-54]. The relationship between HRV (i.e., autonomic function), and glymphatic function during NREM sleep in Veterans with RBD+NT is relevant given the shared connection with neurodegenerative outcomes [55-58]. Impaired glymphatic function has been shown to impair clearance of hallmark neurodegenerative proteins (e.g., amyloid-β, tau, and α-synuclein), with cerebral arterial pulsations driving fluid exchange [48]. RBD is a well-regarded prodromal synucleinopathy, and thus, autonomic impairment via HRV during NREM sleep may also reflect impaired glymphatic function and shed light on a mechanistic determinant of eventual phenoconversion.

It has been proposed by our group and others, that the comorbid condition of RBD+NT is a worthwhile and distinct entity for research [19, 21, 29, 59, 60]. Very few studies have examined HRV with comorbid TBI and PTSD. In a pilot study, Tan et al found the presence of comorbid pain, TBI and PTSD was associated with reduced SDNN during wake, however the sample sizes were small, and the single-condition, clinical groups were not compared to one another [61]. Patients with TBI also have reduced HRV during wake [27, 62-65], however no studies have been conducted examining HRV during sleep. Previous work has demonstrated reduced HRV during wake and sleep in patients with PTSD [66-70]. Aligning with the results of our study, Ulmer and colleagues found Veterans with PTSD had reduced HF power during NREM sleep, indicative of attenuated parasympathetic modulation. They found no differences in HRV during REM sleep [68]. More work needs to be done to untangle the contributing effects of RBD, PTSD, and TBI on sleep HRV. Our data support the comorbid condition of RBD+NT has reduced HRV, and this is consistent with literature that demonstrates reduced HRV in RBD, TBI, and PTSD.

The present study is not without limitations. Our study was conducted at a single site, the Veterans Affairs Portland Health Care System; therefore, all participants were Veterans and consequently, the sample composition was comprised of mostly males. Thus, we were unable to determine whether any of the HRV findings were sex dependent. Furthermore, we did not observe any statistically significant differences between the control and the RBD groups. This may be due to low sample size in the RBD group, or our explicit inclusion of TBI and PTSD. Additionally, this study lacks other corroborating measures of autonomic activity; as such, it cannot be parsed whether changes in parasympathetic versus sympathetic systems are contributing to HRV changes. Finally, this study was conducted as a cross-sectional design; future work would benefit from a longitudinal study design to determine the progression of these HRV changes during sleep. Despite these limitations, our study nevertheless demonstrates reduced HRV during NREM sleep in RBD+NT participants.

The current study found reduced heart rate variability during NREM sleep in individuals with comorbid REM sleep behavior disorder and neurotrauma (TBI+PTSD) compared to controls and RBD alone. Since these patients have been found to have an earlier age of RBD symptom onset, increased symptom severity and generally worsened health and neurological function, heart rate variability (HRV) during sleep may be a clinically relevant biomarker for autonomic dysfunction in this high-risk population. Therefore, measuring HRV during sleep in a prodromal neurodegenerative population (e.g., RBD and NT) could be a potentially promising, early, biosignal to aid in diagnosis and prognosis of the disease course.

## Acknowledgments

The authors would like to express their sincere appreciation and gratitude for the participation of our research participants, VAPORHCS research office staff, and technical support from Jackie L. Gottshall, PhD in the development of related HRV python code. This material is the result of work supported with resources and the use of facilities at the VA Portland Health Care System, with financial support from VA RRD CDA-2 #1K2-RX002947, VA BLRD CDA-2 #IK2-BX002712, DOD #W81XWH-17-1-0423, NIH R34-AG056639, and NIH U19-AG071754. The funders had no role in study design, data collection and analysis, decision to publish, or preparation of the manuscript.

## Disclosure Statement

Financial disclosure: None.

Nonfinancial disclosure: None.

The interpretations and conclusions expressed in this article are those of the authors and do not necessarily reflect the position or policy of the Department of Veterans Affairs, the National Institute of Health, or the United States government.

